# Adaptation to temporally fluctuating environments by the evolution of maternal effects

**DOI:** 10.1101/023044

**Authors:** Snigdhadip Dey, Stephen R. Proulx, Henrique Teotónio

**Affiliations:** Institut de Biologie de l’École Normale Supérieure, INSERM U1024, CNRS UMR 8197, Paris, France; Department of Ecology, Evolution, and Marine Biology, University of California, Santa Barbara, U.S.A.

**Keywords:** experimental evolution, *Caenorhabditis elegans*, anoxia, glycogen, bet-hedging, transgenerational effects, fitness

## Abstract

Most organisms live in ever-challenging temporally fluctuating environments. Theory suggests that the evolution of anticipatory (or deterministic) maternal effects underlies adaptation to environments that regularly fluctuate every other generation because of selection for increased offspring performance. Evolution of maternal bet-hedging reproductive strategies that randomize offspring phenotypes is in turn expected to underlie adaptation to irregularly fluctuating environments. Although maternal effects are ubiquitous their adaptive significance is unknown since they can easily evolve as a correlated response to selection for increased maternal performance. Using the nematode *Caenorhabditis elegans*, we show the experimental evolution of maternal provisioning of offspring with glycogen, in populations facing a novel anoxia hatching environment every other generation. As expected with the evolution of deterministic maternal effects, improved embryo hatching survival under anoxia evolved at the expense of fecundity and glycogen provisioning when mothers experienced anoxia early in life. Unexpectedly, populations facing an irregularly fluctuating anoxia hatching environment failed to evolve maternal bet-hedging reproductive strategies. Instead, adaptation in these populations should have occurred through the evolution of balancing trade-offs over multiple generations, since they evolved reduced fitness over successive generations in anoxia but did not go extinct during experimental evolution. Mathematical modelling confirms our conclusion that adaptation to a wide range of patterns of environmental fluctuations hinges on the existence of deterministic maternal effects, and that they are generally much more likely to contribute to adaptation than maternal bet-hedging reproductive strategies.

## Introduction

In environments that fluctuate in a regular way across generations, such as seasonal climate changes bivoltine insects experience, the maternal environment is a reliable cue for the offspring environment. Natural selection in these circumstances creates covariances between the genes individuals pass on and the environments their mothers have experienced [1,2], which can lead to the evolution of transgenerational plasticity or anticipatory maternal effects [3-6]. Hereafter, we refer to “deterministic” maternal effects when the mother’s phenotype in a given environment has a consistent effect on the offspring phenotype in spite of offspring genotype. Conversely, when the environment fluctuates irregularly across generations, such as the erratic water availability that xeric organisms may face in deserts, the maternal environment is not a reliable cue of the offspring environment. While deterministic maternal effects cannot evolve in this context, bet-hedging reproductive strategies can be favored whereby mothers’ produce offspring with a randomized mix of phenotypes, ensuring that at least some will be able to survive and reproduce no matter which environment they experience [7, 8]. Bet-hedging strategies, here called “randomizing” maternal effects, are expected to be adaptive when they increase long-term growth rates [9-12].

Maternal effects, particularly deterministic maternal effects, are common in a wide variety of fungi, plants and animals [13-18]. It is, however, questionable if maternal effects are adaptive [14,18-21], because they may simply evolve as a correlated response to selection for increased maternal performance [22]. If maternal effects are adaptive in fluctuating environments they must evolve because of selection for increased offspring performance, despite potential selection on maternal traits. The evolution of maternal effects is further expected to be correlated with the evolution of compromised maternal performance [6, 24, 25].

Taking advantage of the *Caenorhabditis elegans* mother-offspring conflict between survival in hyperosmotic and anoxic environments as our paradigm [26], we here show that the evolution of maternal glycogen provisioning underlies experimental adaptation to a novel and regularly fluctuating anoxia environment because of selection for improved embryo hatchability in anoxia. We also demonstrate that the evolution of maternal effects is detrimental to the mothers. While populations experiencing predictable changes in rearing anoxia evolved deterministic maternal effects, populations facing an irregularly fluctuating anoxia environment failed to evolve maternal bet-hedging reproductive strategies, instead relying on the evolution of long-term transgenerational effects for adaptation. Mathematical modelling supports our conclusion that, once evolved, deterministic maternal effects and associated fitness benefits may underlie adaptation to a wide range of temporally fluctuating environments.

## Results and Discussion

### Hyperosmotic maternal survival hinders offspring anoxia survival

When hermaphrodites of the N2 genotype of *C. elegans* experience an osmotic stress their embryos show much reduced hatchability to the first larval stage if facing an anoxia stress [26]. This maternal effect is due to a metabolic trade-off between the hermaphrodites’ ability to produce glycerol during growth from larval stages to maturity and the ability to provision their embryos with glycogen [26,2]. We confirmed for several genotypes derived from our lab-adapted population that hermaphrodites reared in high NaCl concentrations resulted in a survival cost to their anoxia-exposed embryos [28,2] (Fig. S1). We therefore asked if adaptation to fluctuating oxygen levels across mother-offspring generations could be achieved by the evolution of strategic glycogen provisioning.

To address this question, we first evolved our lab-adapted population to high NaCl concentrations [30], so that evolution under fluctuating oxygen levels across mother-offspring generations would not be confounded with adaptation to a hyperosmotic challenge. A single salt-adapted population was then used as the ancestor for all experimental populations in the present study. This ancestor population had appreciable standing genetic diversity although males were present at a low frequency so that most reproduction was done by self-fertilization [30] (Fig. S2). Further, this ancestor population showed a marked reduction in fitness when embryos were exposed to anoxia, measured as the per capita growth rate (Fig. 1A), independently of the maternal hatching environment, indicating that it was well adapted to normoxia but had a substantial potential for selection on embryo hatchability in anoxia (Fig. 1B; Linear mixed effects modelling LMM, followed by planned Tukey t test with corrected Kenward-Roger degrees of freedom: t_93.7_ P<0.001; there was no interaction between offspring and maternal hatching environments).

**Fig. 1.**
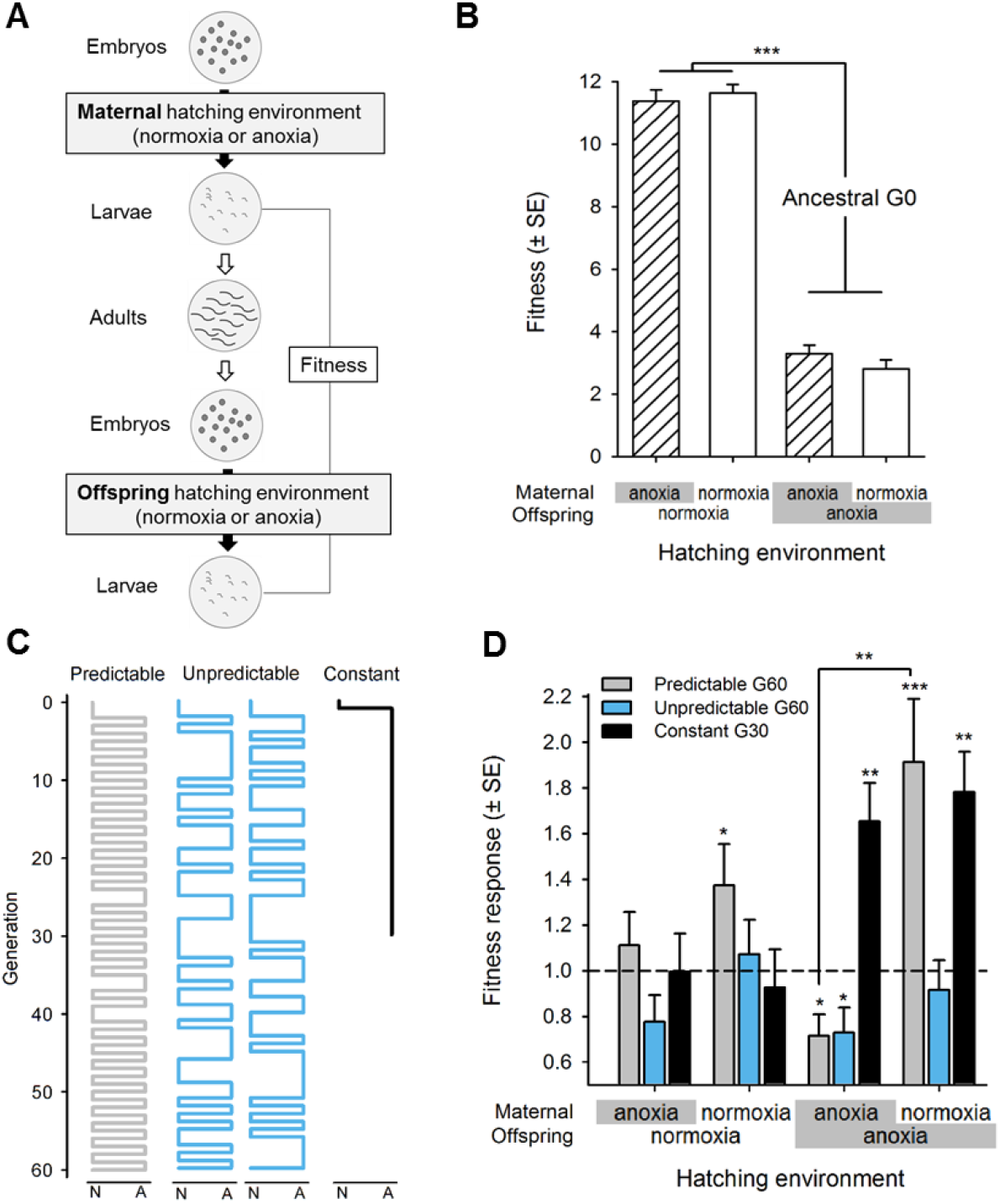
Experimental evolution in temporally fluctuating environments. (A) Four possible mother-offspring environments were implemented during experimental evolution by varying the maternal and offspring hatching environments between normoxia and anoxia. Fitness is defined as the per capita growth rate measured at the first larval stage over one life-cycle. (B) Fitness of the ancestral population across all four maternal-offspring hatching environments. Offspring survival after anoxia exposure is severely hampered, independently of maternal hatching environment, when compared of the normoxia conditions to which the ancestor was adapted (14). (C) Environmental sequences of normoxia (N) and anoxia (A) imposed for 60 generations. (D) After experimental evolution, fitness of predictable (grey), unpredictable (blue) and constant (black) populations relative to the ancestor population (dashed line), across the four combinations of maternal-offspring hatching environments. Mean and error least square estimates are shown linear mixed effects models LMM (B, D). Significant Student t test fitness responses above each bar and planned post-hoc Tukey t test contrasts among hatching treatments: *P<0.05; **P<0.01; ***P<0.001.

### Experimental evolution under fluctuating anoxia hatching environments

Experimental evolution to fluctuating oxygen levels across mother-offspring generations was done for 60 generations under two regimes, each four-fold replicated (Fig. 1C). The “predictable” regime imposed alternating normoxia and anoxia conditions every other generation. Specifically, we designed an environmental sequence where the probability of mothers and offspring sharing the same hatching environment was 0.05, while making sure that the total number of normoxia versus anoxia generations were equal. Similarly, the “unpredictable” regime had an equal number of normoxia and anoxia generations but two environmental sequences were chosen where the probability of mothers and their offspring sharing the same hatching environment was 0.45. This design ensures that non-maternal effect genotypes can perform equally well regardless of the order of the anoxia-normoxia generations in the experiment. A “constant” control regime was characterized by 30 consecutive generations of anoxia, also four-fold replicated. Adaptation in this regime was expected to occur through the evolution of increased embryo hatchability in anoxia, or increased hermaphrodite fecundity following anoxia, regardless of the existence of maternal effects. As was the case during previous laboratory adaptation [28] and previous salt-adaptation [30], all 12 experimental populations were maintained under discrete time and non-overlapping 4 day life-cycles at fixed larval L1 stage to reproductive adulthood density of N=10^4^, such that most adaptation should have occurred from the sorting of standing genetic diversity [31,32].

After experimental evolution, the predictable populations showed a considerable increase in fitness when hermaphrodites experienced normoxia and their embryos experienced anoxia (Fig. 1D; t_15.7_ P<0.001), but showed no change in fitness when hermaphrodites experienced anoxia and their embryos experienced normoxia. Interestingly, there were also changes in fitness in the two mother-offspring environments that were largely absent from the sequences imposed on the predictable populations during experimental evolution. Under two successive generations of normoxia fitness increased (t_10.9_ P=0.03), while under two successive generations of anoxia fitness decreased (t_11.2_ P=0.02). Because these two mother-offspring environmental transitions were not under selection during experimental evolution, they must have evolved as correlated effects.

In contrast to the predictable populations, the unpredictable populations did not improve their fitness in any combination of mother-offspring hatching environments after experimental evolution. In fact, and similarly to the predictable populations, the unpredictable populations showed a significant fitness reduction when exposed to anoxia for two consecutive generations (t_18.5_ P=0.04).

The constant populations evolved increased fitness in anoxia after 30 generations of continuous exposure to anoxia, regardless of the maternal hatching environment (anoxia: t_7.2_ P<0.01; normoxia: t_8.5_ P<0.01). Because our design exposed all experimental populations to the same number of anoxia generations, we can therefore rule out the possibility that lack of genetic variation, small genetic effects, or small population sizes limited the scope for adaptation to anoxia.

### Evolution of deterministic maternal effects trades-off with maternal fecundity

Are these fitness responses of the predictable populations consistent with the evolution of deterministic maternal effects? There are two components of life-history that could be involved in adaptation to fluctuating anoxia: changes in the fecundity of adult hermaphrodites (a direct effect) and changes in embryo hatchability due to maternal glycogen provisioning (a deterministic maternal effect).

In the predictable populations, we found that adult hermaphrodites who experienced anoxia early in life had reduced fecundity (Fig. 2A; t_3.6_ P=0.03), and those that experienced normoxia had similar fecundity (t_3.9_ P=0.4), when compared to ancestral hermaphrodites. Remarkably, a clear deterministic maternal effect is seen in the hatchability response in that hermaphrodites exposed to normoxia early in life produce embryos with increased hatchability in general, with a larger response under anoxia (Fig. 2B; offspring anoxia: t_14.5_ P<0.01, offspring normoxia: t_12.9_ P=0.03). These results thus parallel the fitness results and provide an explanation for the observed correlated response of an increase in normoxia-normoxia fitness (an environmental sequence that the predictable populations did not face during experimental evolution). Specifically, evolution of increased embryo hatchability in normoxia resulted in increased fitness in normoxia. These results additionally demonstrate the evolution of a trade-off between embryo survival under anoxia and adult hermaphrodite fecundity after hatching under anoxia, in line with theoretical expectations [6,24,25].

**Fig. 2.**
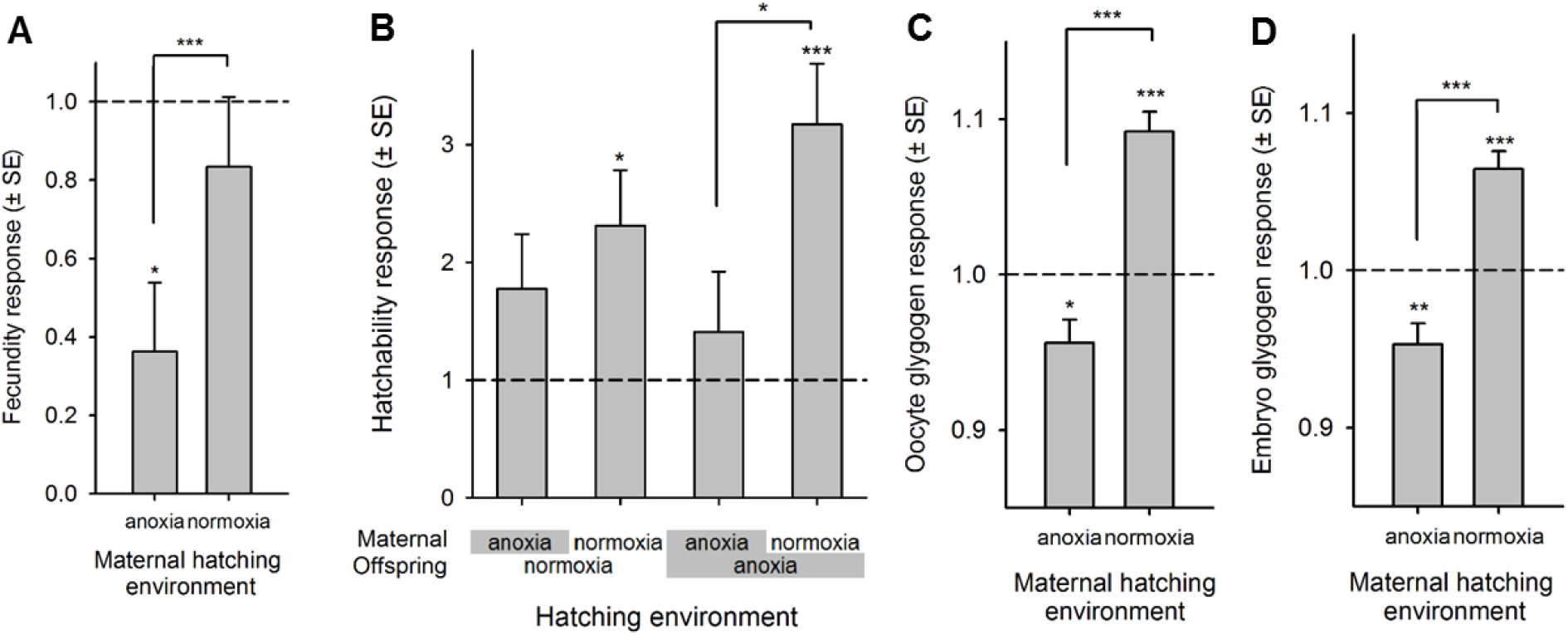
Experimental evolution of maternal glycogen provisioning protects embryos from anoxia but at the expense of fecundity. Fecundity (A), hatchability (B), oocyte (C) and embryo (D) glycogen content of the predictable populations after experimental evolution relative to the ancestral population (dashed line). For all panels, mean and error least square estimates are shown after LMM, significant evolutionary responses above each bar, and planned Tukey t test contrasts among oxygen level treatments: *P<0.05; **P<0.01; ***P<0.001.

### Evolution of maternal provisioning of glycogen to offspring

We directly confirmed the hypothesis that the deterministic maternal effect that evolved in the predictable populations was mediated by changes in strategic maternal glycogen provisioning [26]. Image analysis of iodine-stained oocytes and early embryos within hermaphrodites showed the evolution of increased glycogen content only when they experienced normoxia early in life (Fig. 2CD; oocytes: t_8.8_ P<0.01, embryos: t_7.6_ P<0.001; see also Fig. S3). Surprisingly, both oocyte and embryo glycogen content was reduced relative to the ancestor population in anoxia-exposed hermaphrodites (oocytes: t_8.2_ P=0.03, embryos: t_11.3_ P<0.01), suggesting, like fecundity, a trade-off between embryo anoxia survival and hermaphrodite glycogen provisioning after being reared under anoxia. The evolution of reduced glycogen provisioning appears to be inconsequential, however, since embryo hatchability in normoxia --the environment the embryos produced by hermaphrodites hatched under anoxia would normally face--did not change during experimental evolution.

### Adaptation to irregularly fluctuating environments does not involve maternal bet-hedging strategies

Having shown the evolution of a deterministic maternal effect in the predictable populations, we address whether there was evolution of a randomizing maternal effect in the unpredictable populations. In temporally fluctuating environments the appropriate measure of adaptation is the geometric mean of environment-specific fitness [9,10,12]. The presence of maternal effects (deterministic or randomizing) does not change this principle, but does require averaging over the frequencies of the pairwise environmental transitions experienced by the population during its history. With uncorrelated environmental variation, reproductive strategies that reduce generation-to-generation variation in maternal performance and increase the geometric mean of fitness are known as bet-hedging strategies [7,8,18]. If there was evolution of maternal bet-hedging in the unpredictable populations then their geometric mean fitness, averaged over the four pairwise mother-offspring hatching environments, should have improved relative to the ancestor population.

We used a two-generation model of maternal effect carryover to estimate the geometric mean fitness of the unpredictable populations and found no evidence that it changed with experimental evolution (Fig. 3A; t_6_ P=0.19). The predictable populations, as expected, increased their geometric mean fitness across the two environmental transitions they experienced during their history (t_6_ P<0.01). Numerical simulations indicate that this lack of response in the unpredictable populations is not due to the fact that we measured the geometric mean fitness of genetically variable populations [8,11] (Fig. 3BC; Fig. S4). Further, since unpredictable populations did not go extinct during experimental evolution, another component of the long-term fitness must have increased enough to compensate the observed reduction in anoxia survival over two successive generations (vid. Fig. 1D). Multi-generational carryover effects are indeed expected to be adaptive in irregularly fluctuating environments [25,33-36], and there was scope for their evolution since the unpredictable populations faced a set of environmental transitions that were correlated over the long-term.

**Fig. 3.**
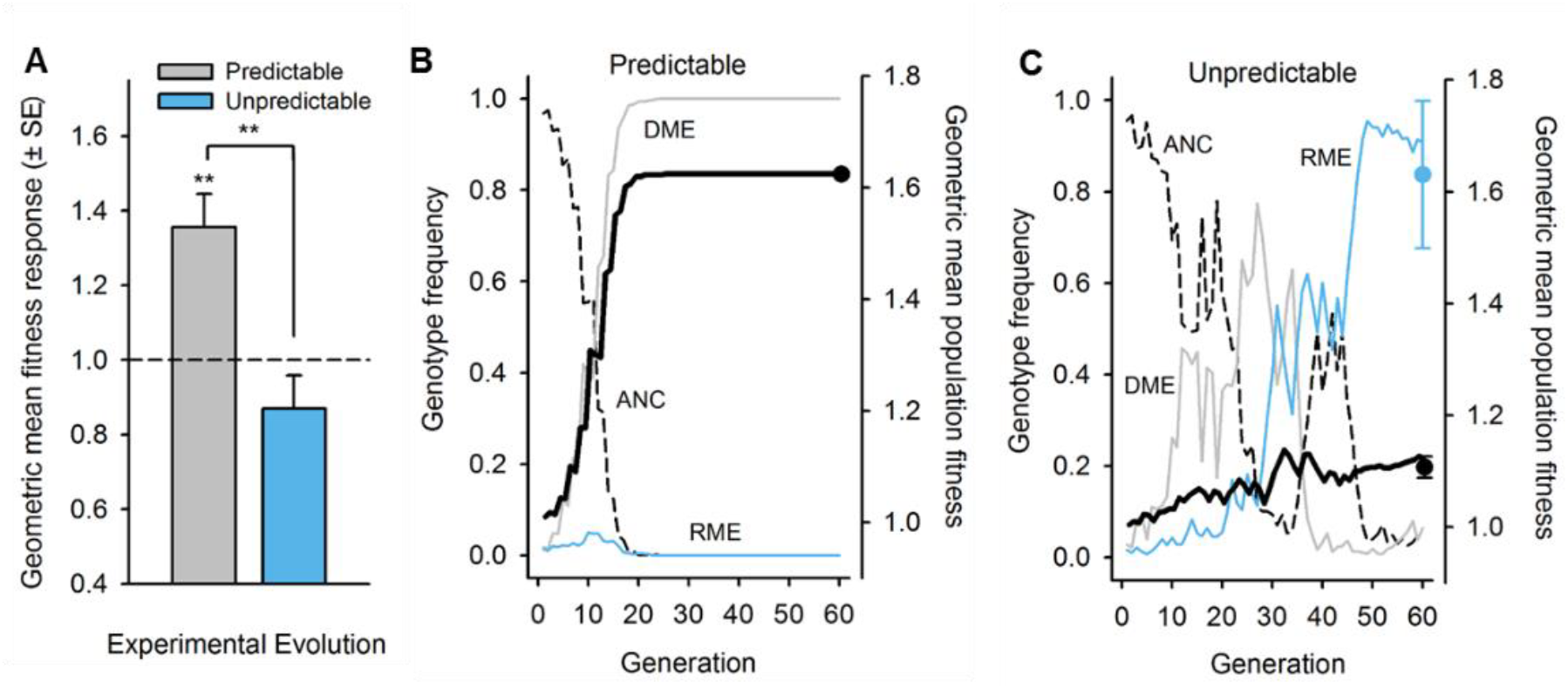
Maternal bet-hedging does not explain adaptation under irregular anoxia environments. (A) The geometric mean fitness of the oxygen level transitions across generations experienced by the predictable (normoxia-anoxia and anoxia-normoxia) and unpredictable populations (all four pairwise combinations) relative to the ancestral state (dashed line). Mean and error least square estimates are shown after ANOVA, significant evolutionary responses above each bar, and an F_1,6_ test contrast among experimental regimes: **P<0.01. (B, C) Simulations mimicking the evolution of maternal effects during experimental evolution (see Methods). (B) The probability of environmental transition was set to 0.95, as the predictable populations faced during experimental evolution. Relative phenotypic fitness values are of W_A-to-N_=0.335 and W_A-to-A_=2.63, similar to those of Fig. 1D. The deterministic maternal effect genotype (DME; grey line) reaches fixation at generation 24, during which the population geometric mean fitness (black line) smoothly increased to the expected value of the square root of 2.63 = 1.624. Fifty simulations with the same parameter values show that at generation 60 there is always fixation of the DME genotype and thus the same population geometric mean fitness (black circle). The randomized maternal effect genotype (RME; blue line) does not invade the population composed by the ancestral genotype that does not express maternal effects (ANC; dashed line). (C) As in panel (B) but with the probability of environmental change at 0.55.

### Which maternal effects underlie adaptation to fluctuating environments?

To better understand how the sequence of environmental transitions determines the course of evolution we developed mathematical models to calculate the probability that a genotype conferring maternal effects would invade the ancestral population [11,37] (see Methods). We considered a scenario with two offspring phenotypes, one that had higher fitness under anoxia and the other that had higher fitness under normoxia. Deterministic maternal effects were defined by a genotype that always produced offspring with the phenotype suited to the environment that the mother did not experience. The randomized maternal effects genotype produced a brood of offspring with a fixed fraction of each phenotype, with the fraction chosen in order to maximize the geometric mean fitness of the bet-hedging strategy.

We found that the deterministic maternal effect can only evolve if the probability of switching environments is above 0.5, and increases as the probability of switching goes up (Fig. 4A). The maternal bet-hedging strategy has a constant benefit relative to the ancestor for switching rates of less than 0.5, and is outcompeted by the deterministic maternal effect strategy when the probability of switching is large. The maximum probability of fixation for the randomizing bet-hedging strategy is only about 1/10 that of the deterministic maternal effect, even if the same environment-specific phenotypic effects were parameterized in both kinds of maternal effects.

**Fig. 4.**
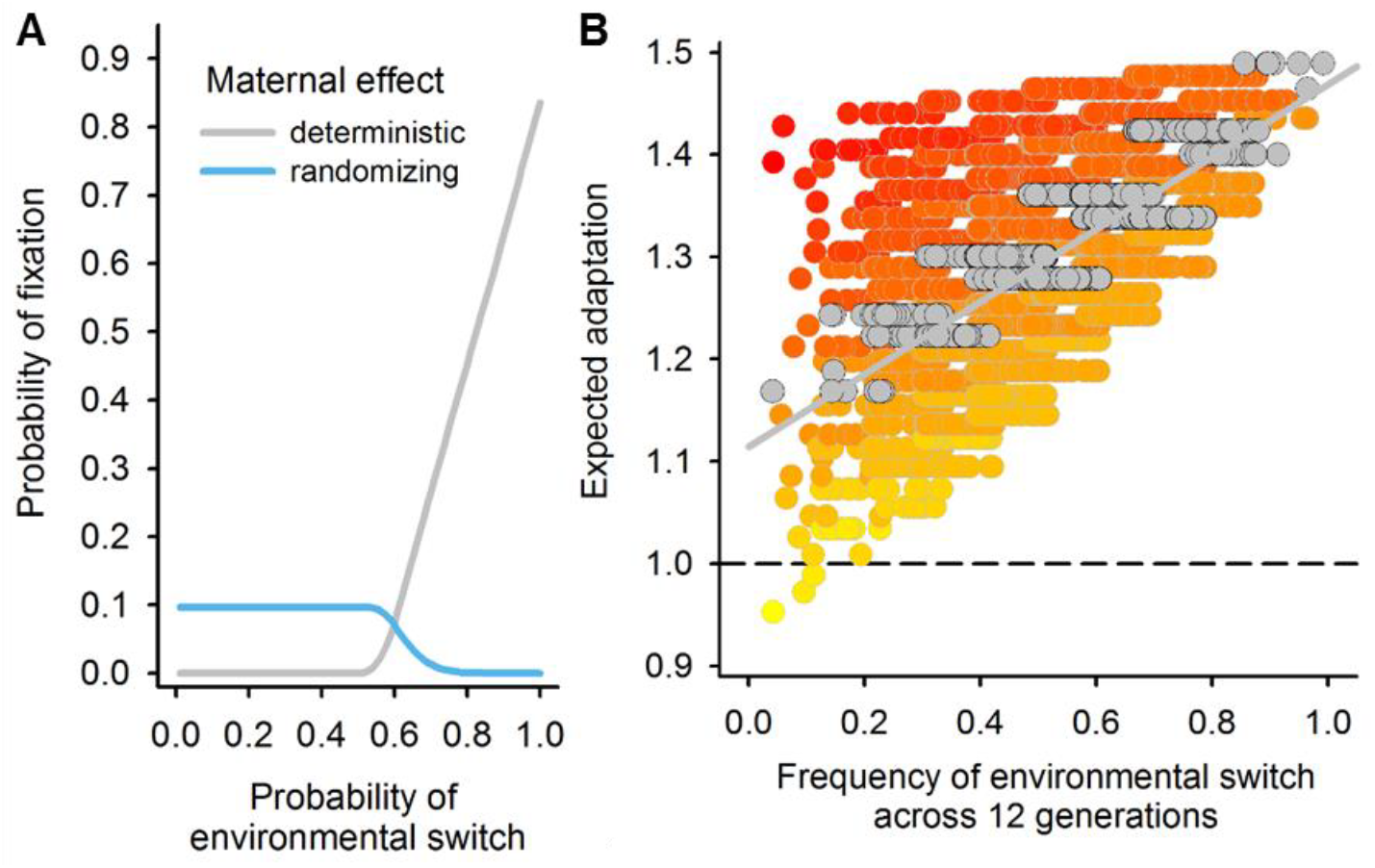
Maternal deterministic effects and associated fitness benefits can underlie adaptation to fluctuating environments. (A) The probability of fixation of a genotype expressing deterministic maternal effects (gray line) or randomized maternal effects (blue line), when invading the ancestral population without maternal effects. Results are from 10,000 randomly drawn fitness parameters with *γ* = 2.0, producing approximately 2-fold fitness effects on average (from Fig. 1D). Other distributions yield qualitatively similar results. Fixation probabilities were calculated with effective population size of 10^3^ [32,2], and an initial frequency of 0.01 (see Methods). (B) Expected adaptation of the predictable populations relative to the ancestor population (dashed line) if they were to face fluctuating sequences of normoxia and anoxia for 12 generations. Data is jittered along the x-axis for clarity. Each circle shows one environmental sequence out of the 2^12^ possible, with the yellow-to-red color gradient depicting increasing frequency of normoxia generations, and grey circles and line representing sequences in which normoxia and anoxia generations are equally represented.

Experimental evolution results are thus consistent with our theoretical analysis, particularly because adaptation caused by maternal effects was only observed in the predictable populations. To illustrate that deterministic maternal effects are potentially adaptive in a range of patterns of environmental fluctuations, we used the evolved predictable populations’ two-generation fitness values to calculate the geometric mean fitness for arbitrary sequences of fluctuating oxygen level hatching environments (Fig. 4B). The simulations show that the evolved genotypes would perform better than the ancestral genotypes even in environmental fluctuation regimes that were not used in experimental evolution. This suggests that the evolution of deterministic maternal effects and correlated fitness benefits unlocked an adaptive potential that was not accessible to the ancestral population. We conclude that adaptation to regularly fluctuating environments may improve a populations’ future prospects of withstanding a wide array of environmental fluctuation patterns.

## Conclusion

Our analysis demonstrates that deterministic maternal effects, and possibly multi-generational carryover effects, can underlie adaptation to temporally fluctuating environments. We find much less support that maternal bet-hedging reproductive strategies randomizing offspring phenotypes make a strong contribution to adaptation. With anthropogenic activities increasingly contributing to more extreme and irregular climate fluctuations [38], it will be important to understand if deterministic maternal effects are also key in maintaining biodiversity and in preventing extinction [39].

## Materials and Methods

### Salt-anoxia survivorship in the lab-adapted population

Four inbred lines of a lab-adapted population, A6140, were derived by 12 generations of enforced self-fertilization as previously described [29] (A6140L126, A6140L142, A6140L188, A6140L244), and stocks frozen at -80°C. On the third generation after thawing and passaging under the standard lab conditions, first stage larvae (L1s) from each of the inbred lines were seeded in either 25mM NaCl or 300mM NaCl standard NGM-lite plates with *E. coli* [28]. After 96h, fifty embryos were hand-picked and transferred to 6cm Petri dishes with 25mM NaCl NGM-lite and a 10uL drop of *E. coli*, and then placed under normoxia or anoxia conditions for 16h (see below). After 96h, the number of adults was scored. For each inbred line and maternal-offspring treatment, 5 replicates were done. For statistical analysis, ANOVA was done with inbred line (4 levels) and maternal-offspring treatment (4 levels) as fixed factors. F-tests were employed to test for significance of main factors and interaction. Post-hoc Tukey t-tests were then done to contrast maternal-offspring treatments. In a separate assay, line A6140L244 was similarly assayed for anoxia survivorship from embryo to adulthood, when hermaphrodites from the maternal generation were reared from 24h to 96h of the life-cycle in 25mM, 100mM, 200mM, 225mM, 250mM, 275mM and 300mM NaCl NGM-lite plates. Twenty replicates per treatment were done.

### Environmental sequences

The predictable environmental sequence of anoxia-normoxia generations was designed such that the probability of repeating the same oxygen level hatching conditions across two generations was of 0.05, and the frequency of anoxia and normoxia events was 0.5 for a total of 60 generations. The two unpredictable sequences (#11, #19) were designed such that the probability of repeating the same oxygen level hatching conditions across two generations was of 0.45 and across three generations of 0.5. The frequency of anoxia and normoxia generations was 0.5 for a total of 60 generations. The constant environment was characterized by 30 consecutive generations of anoxia. QBasic64 v0.954 was used to design the sequences.

### Experimental evolution

All populations were derived from a high salt-adapted population [30] (GA250), which in turn was derived from a lab-adapted population [28,29] (A6140). Frozen GA250 samples with >10^4^ individuals were thawed from frozen -80°C stocks and cultured for one generation for number expansion, before replicate population derivation. Predictable, unpredictable and constant populations were designated Pi, Ui and Ci, respectively, with *i* standing for 1 to 4 replicate number. During experimental evolution a random number was assigned to each population to avoid potential bias. Environmental sequences #11 and #19 were each replicated in two populations. Following our standard laboratory environment [28,30], populations were kept in ten 9 cm Petri plates with 28mL of solid NGM-lite agar media (Europe Bioproducts) covered by an overnight grown lawn of HT115 *E. coli* food. NaCl concentration in the NGM-lite media was of 305mM (1.78% w/v)[30]. At 24h±2h of the life cycle, each population was seeded with 1,000 first larval staged (L1) individuals in each of the ten Petri plates. After growth to maturity for 66h±1h at constant 20°C and 80%RH in controlled incubators (Fitoclima D1200, ARALAB), all ten plates were mixed and worms harvested with 5mL M9 isotonic solution and then exposed to 1M KOH: 5% NaOCl “bleach” solution for 5min±15sec, to which only embryos survive. After repeated washes with M9, 200μL containing embryos (and larval and adult debris) were transferred to 25mM NaCl NGM-lite plates, without *E. coli*. These plates were then placed inside 7L polycarbonate boxes with rubber clamp sealed lids (AnaeroPack, Mitsubishi Inc.). Within these boxes an anoxic embryo hatching condition was imposed by placing two GasPak^TM^ EZ sachets (Becton, Dickinson and Company). These sachets contain inorganic carbonate, activated carbon, ascorbic acid and water, which after two hours of activation will produce an anaerobic/anoxic atmosphere inside the boxes (<1% O_2_, ≥13% CO_2_; according to the manufacturer). At every generation, in all boxes, anoxia conditions were confirmed by placing two BBL^TM^ Dry Anaerobic Indicator strips (Becton, Dickinson and Company). To prevent drying, paper towels with 20mL ddH_2_0 were placed within each box. For normoxia hatching conditions, the sachets were not used and an additional 60mL ddH20 was placed inside each box. After 16h, hatched L1s were washed off the plates with 3-5mL M9 to a 15mL Falcon tube, adult debris removed after centrifugation at 200rpm and, live L1 density estimated under a Nikon SMZ1500 dissection scope in five 5uL M9 drops at 40x magnification. While estimating, tubes were kept in a shaker at 120rpm (Lab. Companion SK-600) inside the incubators. 6h±1h after, the appropriate M9 volume for 1,000 live L1s was placed in fresh NGM-lite plates to complete one life cycle.

### Fitness assays

P1-4 and U1-4 from generation 60 and C1-4 from generation 30 were measured for fitness alongside the ancestral GA250 population in two-generation long assays. Frozen -80°C stocks (n>10^3^) were thawed and reared in a common environment for two generations before assaying. On the third generation, adults were washed off the plates, treated with the “bleach” solution and their embryos were exposed to normoxia or anoxia to constitute the maternal hatching assay environment. 24h later, 1,000 of surviving L1s were seeded in each of 5 Petri dish plates, allowed to grow to adulthood, treated with the “bleach” solution and their embryos were exposed in a factorial fashion to normoxia or anoxia to constitute the offspring hatching assay environment. After 16h of exposure to the corresponding hatching environment, worms were washed off the plates with 3-5mL of M9 to a 15mL Falcon tube before the total number of surviving L1s was estimated by considering the total volume of the M9 solution and counting the number of live L1s in ten to fifteen 5μL drops. The total number of live L1s was then divided by 5,000 to calculate the maternal L1 to offspring L1 growth rate. This ratio defined fitness in our two-generation paradigm as the per capita growth rate. The assays were done in 16 blocks, defined by different thawing dates of the samples. In each block the ancestral GA250 population was included. Per population and hatching environmental treatments there were 3 replicate measurements.

### Fecundity and hatchability assays

These assays were similar in design to the fitness assays, with two-generation exposure to all four mother-offspring oxygen hatching environment combinations. P1-4 from generation 60 and the ancestor GA250 (n>10^3^) were thawed and grown in parallel for two generations before assaying. For each replicate measurement, 1,000 live L1s after exposure to normoxia or anoxia maternal hatching environments were grown in 6 to 10 plates at a density of 1,000. Adult worms were washed off and treated with the “bleach” solution. In contrast with the fitness assays, after the “bleach” treatment the dead adults were removed from the M9 Falcon tube after centrifugation at 200rpm. The number of embryos was then scored in ten 5μL M9 drops to estimate the total number of embryos in the M9 volume after the “bleach” treatment. This total number was then divided by the total number of adults to calculate the per capita fecundity. The embryo-only M9 solution was equally divided within 2h after the “bleach” treatment, centrifuged at 1,800rpm. The pellet containing the embryos was then exposed to anoxia or normoxia hatching conditions. After 16h the density of live L1s was estimated. Hatchability was then calculated as the ratio of live L1s over embryo number. The assays were done in two thawing time blocks, for 3 replicate measurements per population and hatching condition.

### Glycogen content assays

Adult hermaphrodites from generation 60 P1-4 and GA250 populations were assayed for glycogen content in the oocytes and embryos inside the body. After exposure to a normoxia or an anoxia hatching environment, adult hermaphrodites were subjected to fixation. This was done by suspending them in a solution of 700 μL of absolute ethanol (VWR Scientific) with 200 μL of glacial acetic acid (Carlo Erba) and 100 μl of concentrated formalin (Sigma-Aldrich) for 90 minutes. Serial dehydration was then done by sequentially re-suspending worms in 70%, 90% and absolute ethanol. Samples were stored at -20°C. Frozen hermaphrodites were sequentially rehydrated in absolute, 90% and 70% ethanol, before being re-suspended in the M9 solution. Hermaphrodites were then transferred to a glass slide topped with a thin agar pad (5% noble agar; Becton, Dickinson and Company) with a pipette. Hermaphrodites from the ancestral and one of the predictable populations were assayed simultaneously by placing them on the same agar pad. For staining, each glass slide was placed upside down over the mouth of a bottle with 100g of iodine (I_2_, ACS≥99.8% solid; Sigma-Aldrich) for 120 seconds [26,27]. Within 40 minutes, photographs were taken at 630x magnification under differential interference contrast (DIC) settings in a Zeiss Axioskop2 microscope coupled to a monochromatic CCD camera (Hitachi Denshi ltd.). The DIC settings for the microscope and camera were kept identical across all glass slides. For each hermaphrodite, 2-3 images were taken to cover developing embryos and/or oocytes at the anterior half of the body from the vulval region. Similar images were obtained for unstained samples. The images were not manipulated for contrast of grayscale levels. For analysis we used ImageJ 1.46r, with the first three oocytes from the spermatheca, or all visible embryos, being manually delineated and the mean pixel intensity over the measured area recorded. For all the images, the mean pixel intensity of agar pad was also obtained to account for the decay of staining with time and/or other non-specific staining variation across images. The ratio of the mean pixel intensity of the oocytes/embryos over the mean pixel intensity of the agar pad was used as the raw data for statistical analysis. Size was taken as the perimeter in pixels of the individually delineated oocytes. The assays were run in one block, for 3 to 8 replicate measurements per population and hatching oxygen condition.

### Statistical data analysis

Fitness data in the ancestral population was analyzed with a linear mixed effects model (LMM) and REML estimation methods [40], taking maternal and offspring hatching conditions as fixed two level factors, and block as a random factor. A preliminary model revealed that residuals did not follow normality (as tested with the Shapiro-Wilk test), and thus outliers were removed through the inspection of Quantile-Quantile Plots. Contrasts were then performed among the mother-offspring hatching conditions, using post-hoc Tukey t-tests while estimating the effective number of degrees of freedom with the Kenward-Roger (KR) approximation. For evolutionary responses, we transformed the data from the derived populations by taking the ratio to the ancestral mean value per block. Transformed data was analyzed by LMM, taking replicate population as the random factor in order to account for the effects of genetic drift and other historical accidents. Fitness response data was log-transformed to solve normality issues, with the least-square estimates presented being the back-transforms. As above, the KR approximation was used to estimate degrees of freedom in planned Tukey t-tests. For fitness responses, we first modelled experimental evolution regime (2 levels, predictable and unpredictable) and maternal-offspring hatching conditions (4 levels) as the fixed factors. Generation 30 fitness data from the constant populations was analyzed separately since the extent of genetic drift in them was halved relative to predictable/unpredictable populations. Including the constant populations in the first model would underestimate the random replicate population effects. For fecundity, glycogen content and oocyte size responses, we modelled the maternal hatching environment (two levels) and for the hatchability responses maternal-offspring hatching environments (with four levels) as the fixed factors. For the geometric mean fitness responses, fitness values in normoxia-anoxia and in anoxia-normoxia transitions in predictable populations and in all four pairwise transitions for unpredictable populations were modelled together with ancestral fitness values by LMM, taking block as a random factor. The geometric mean of the estimated mean least-squares were then calculated, with predictable and unpredictable responses taken as the ratio to the ancestral in an ANOVA with experimental regime as a fixed factor. For all models, significant evolutionary responses were inferred when the estimated effects deviated from one (the fixed ancestral state), as tested with Student t-tests and, if needed, KR estimated degrees of freedom. Least-square mean and error estimates are presented in all plots. The packages *stats*, *lme4*, *lsmeans* and *pkbrtest* in R were used for computation [41].

### Genotyping and genetic diversity analysis

Immature hermaphrodites were handpicked from GA250 and generation 60 U1-4 populations and their gDNA collected using prepGEM Insect kit (ZyGEM). Bi-allelic single-nucleotide polymorphisms (SNPs) were chosen based on the pooled genome sequence of the A6140 population (unpublished data). SNPs were genotyped with the iPlex Sequenom^TM^ technology [28,42]. WS200 genome version was used for the oligonucleotide design and SNP physical position (www.wormbase.org). SNPs in chromosome I and II were genotyped in 48 GA250 individuals and in 16 individuals from each of the U1-4 populations, chromosomes III-IV and separately V-VI were genotyped with similar sample sizes. SNPs for which the assay failed in more than 30% of the samples were eliminated. This was followed by elimination of individuals with more than 10% of undetermined allele identity, and finally elimination of SNPs with failed assays in more than 15% of the remaining individuals across populations. A total of 478 SNPs were used for analysis (chr. I, 4.5 SNPs/Mbp; chr.II, 6.1 SNPs/Mbp; chr. III, 6 SNPs/Mbp; chr. IV, 4 SNPs/Mbp; chr. V, 4.8 SNPs/Mbp; chr. VI, 6.9 SNPs/Mbp). Within-population fixation indices (Fis) were calculated as 1 minus the ratio of observed (Ho) to expected heterozygosity levels under random mating [42].

### Mathematical modelling

We took two modelling approaches, one based on a general model of maternal effects that we used to predict when maternal effects are likely to evolve andn other based on the empirically observed fitness responses that we used to predict how the evolved populations would respond to novel environmental sequences. All computations were done using Wolfram Mathematica 9.

In the general model we calculated the geometric mean fitness of genotype *k* as *G*_*k*_ = exp (Σ *p*(*i*, *j*))log (*w*_*k*_ (*i*, *j*))), where *G*_*k*_ represents the geometric mean fitness of genotype *k*, *p*(*i*, *j*) is the frequency of transitions from hatching environment *i* to hatching environment *j*, and *w*_*k*_ (*i*, *j*) is the reproductive output of a genotype *k* individual developing in environment *j* whose mother experienced hatching environment *i*. We model maternal effects by assuming that there are two possible offspring phenotypic states, normoxia adapted and anoxia adapted. Without loss of generality, we assigned the normoxia adapted phenotype relative fitness values of 1, so that *w*_*N*_(*i*, *j*) = 1. We assigned the log fitness of the other anoxia phenotype by drawing from an exponential distribution with parameter *γ* so that fitness in anoxia was increased and fitness in normoxia was decreased. A value of *γ* = 2.0 means that the average advantage of the anoxia adapted phenotype in anoxia conditions is about 2 fold, on the order of the fitness increase we observed following experimental evolution. We assumed for simplicity but without loss of generality that the ancestral phenotype, the normoxia adapted phenotype, had higher geometric mean fitness than the anoxic phenotype.

We then used the geometric mean to calculate the probability of fixation of either a genotype conferring a Deterministic Maternal Effect (DME) or a Randomizing Maternal Effect (RME). The ancestral state was assumed to have no maternal effect and therefore produce a constant phenotype adapted to normoxia conditions. Under DME, mothers alter their offspring phenotype based on their own hatching environment. Under RME, mothers produce a fraction *q* of offspring with the anoxia adapted phenotype and *(1-q)* with the normoxia phenotype. We assumed *q* was tuned to maximize the geometric mean fitness of the genotype, an assumption that favors RME. Given that experimental populations were maintained in discrete time non-overlapping generations, the probability of fixation of an invading genotype (labelled *U*_*DME*_ and *U*_*RME*_, respectively) was calculated using M. Kimura’s approximation of the Wright-Fisher process [11,43]. In line with other evolution experiments from standing genetic diversity, we further assumed that effective populations sizes were one order of magnitude lower than the experimental census sizes [32,42]. We defined the effective selection coefficient for use in Kimura’s equation as the geometric mean of the reproductive output of a genotype, which is expected to be a good approximation so long as the probability a genotype goes extinct due to selection is not abnormally high in early generations [37,44]. Because our populations had standing genetic diversity we wished to calculate the probability that DME or RME would become common based on both strategies being present at some low initial frequency. We therefore adjusted our calculations to include the chance that both strategies would become established at appreciable frequencies, in which case the strategy with higher geometric mean fitness is expected to prevail. Thus, for parameter values that give DME a higher geometric mean this is simply *U*_*DME*_. However, if RME had a higher geometric mean this becomes *U*_*DME*_(1–*U*_*RME*_).

We also performed stochastic simulations of the spread of DME and RME genotypes into an ancestral population composed mostly of genotypes that produce only the normoxia adapted phenotype. We used population size of 1,000 and an initial frequency of both ME strategies of 0.005. RME strategies were assumed to use the optimal frequency of producing normoxia/anoxia adapted phenotypes. Again, we used the Wright-Fisher model to simulate changes in genotype frequency. In each generation the relative number of offspring produced by each genotype was simply the frequency of genotype *i* multiplied by the average fitness of genotype *i* in the current environment. The number of individuals of genotype *i* was updated by sampling from a multinomial distribution where the probability of drawing genotype *i* was the reproductive output of genotype *i* divided by the total reproductive output of the population. We calculated the population level fitnesses as they would have been measured in our experimental assays and used these to calculate the geometric mean fitness of the population.

For our second modelling approach we constructed lists of sequences of 12 generations of normoxia or anoxia and then used the assayed fitness values to calculate the “expected adaptation” of the derived predictable populations relative to the ancestral population. The total frequency change of a genotype over multiple generations is a simple function of the product of the relative fitness of the two genotypes in each generation. We therefore defined the expected adaptation as 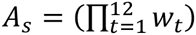, where the relative fitness of the novel genotype is *w*_*t*_ in generation *t*, and *A*_*s*_ measures the per-generation fitness advantage of the novel genotype in an environmental regime characterized by sequence *s*.

## Acknowledgements

We thank S. Carvalho, I.M. Chelo, M.A. Félix, T. Guzella, P. Ibañez, S. Nunes and A. Pino for technical support, and I.M. Chelo, M.A. Félix, T. Guzella, and P.C. Phillips for discussion. S.D. did experimental evolution and assays; S.D. and H.T. analyzed the data; S.R.P. did the modelling; S.D., S.R.P. and H.T. designed the project and wrote the manuscript. Funding provided by the National Science Foundation (EF-1137835) to S.R.P., and the European Research Council (FP7/2007-2013/243285) to H.T.

## Supplementary Figures

**Fig. S1.**
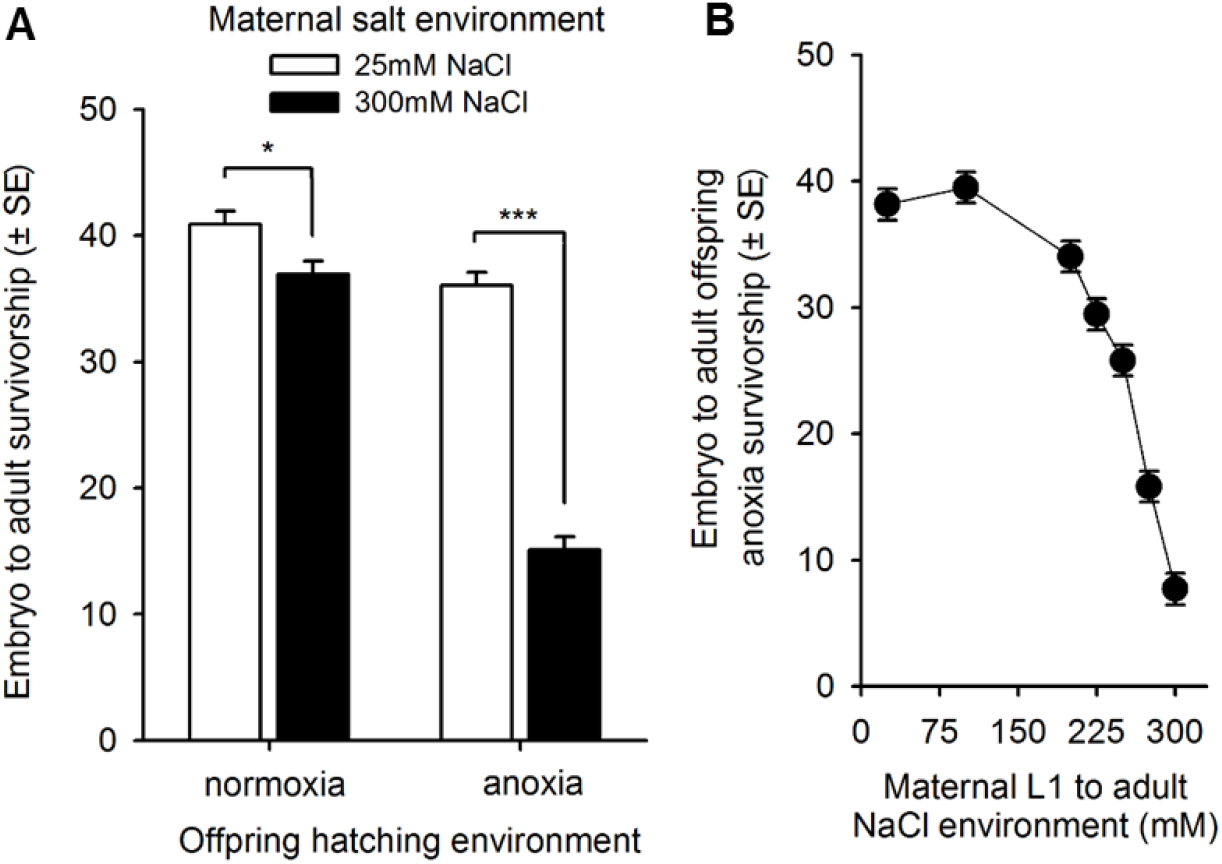
High salt maternal environments impair embryo survival to anoxia. (A) Embryo to adult survivorship under normoxia and anoxia hatching environments, after maternal rearing from L1 larval stage to adulthood at low (white) or high (black) salt concentrations. Four different genotypes (L126, L142, L188, L244) derived from a lab-adapted population (A6140) were used in these assays (14). L1s from each of the inbred lines were seeded in either 25mM NaCl or 300mM NaCl NGM-lite plates with *E. coli*. After 96h, fifty embryos were hand-picked and transferred to 6cm Petri dishes with 25mM NaCl NGM-lite and a 10uL drop of E. coli, and then placed under normoxia or anoxia conditions for 16h (see Materials and Methods). After 96h, the number of adults was scored for survivorship. ANOVA mean and error least square estimates are presented: maternal-offspring treatment: F_3,63_ P<0.001, interaction genotype with maternal-offspring treatment: F_9,63_ P<0.001. Post-hoc Tukey t-tests contrasts among maternal salt treatments: *P<0.05; ***P<0.001. (B) Reaction norm of embryo to adult survivorship over different maternal salt concentration environments for genotype L244 (n=20 per treatment).

**Fig. S2.**
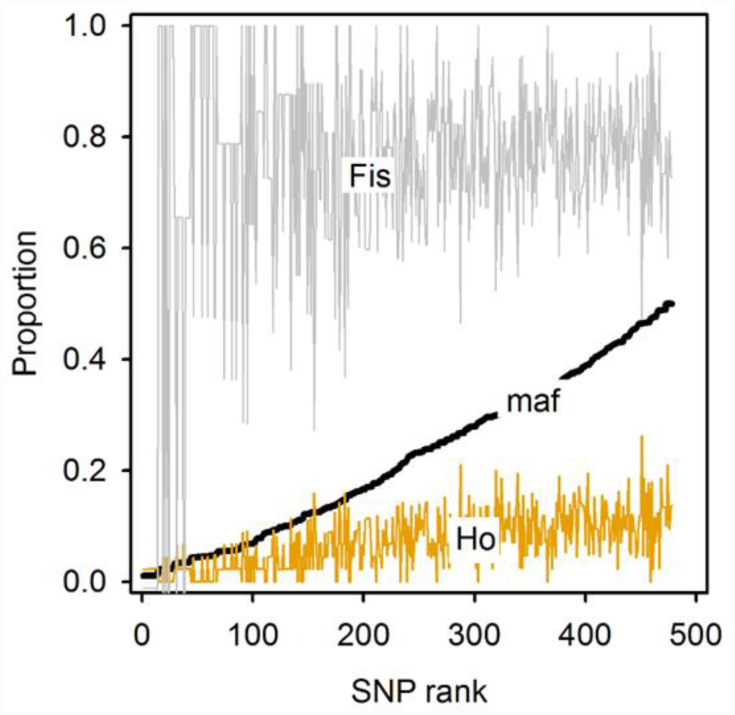
Standing genetic diversity in the high-salt adapted ancestor population. Observed heterozygosity (Ho) and fixation indices (Fis) of 478 SNPs covering the whole genome and ranked according to increasing minor allele frequency (maf). High Fis values indicate a high level of inbreeding due to a high proportion of hermaphrodites reproducing by self-fertilization.

**Fig. S3.**
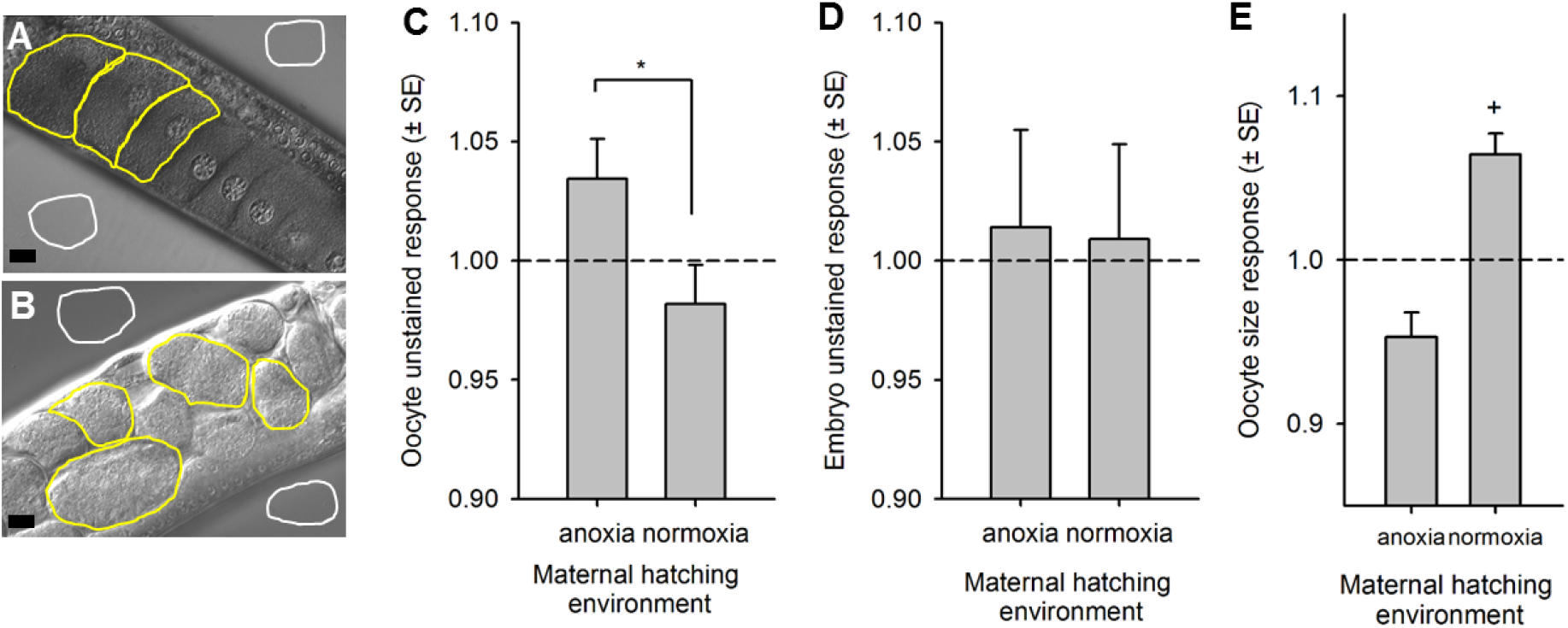
Iodine staining of glycogen in oocytes and embryos. (A, B) Glycogen content was quantified in iodine vapor stained oocytes and embryos within hermaphrodites (see Materials and Methods). Illustrative photographs of stained oocytes and unstained embryos are shown from ancestral hermaphrodites. The width of the scale bar is 10μm. The ratio of the mean pixel intensity of all the delineated oocytes/embryos in an individual (yellow lines) and the mean pixel intensity of the corresponding agar pad was used for analysis (solid white lines). The ratio values of control unstained oocyte (C) and embryo (D), or oocyte size (E), of the predictable populations at generation 60 relative to the ancestor state (dashed line). After LMM, we did not detect evolutionary responses, although values were higher when hatching under anoxia relative to hatching under normoxia (t_24.4_ P=0.01). Oocyte size in normoxia-hatched hermaphrodites may have increased relative to the ancestor (t_9_ P=0.06), but there is no correlation between oocyte size and its pixel intensity ratio in stained hermaphrodites (Pearson’s ρ=-0.04, t_78_ P=0.73).

**Fig. S4.**
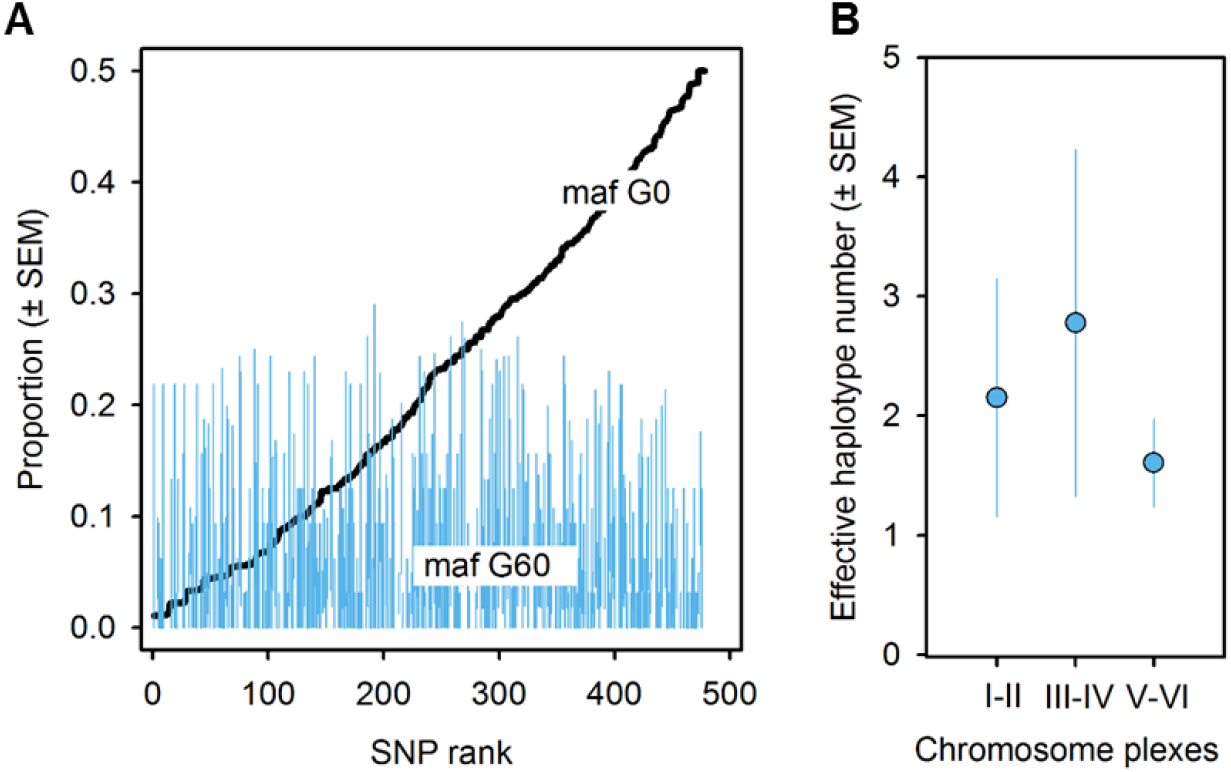
Genetic diversity in the unpredictable populations. (A) 478 SNPs are ranked according to increased minor allele frequencies in the ancestor population (from Fig. S2, G0), and compared to the minor allele frequencies of the unpredictable populations at generation 60 (G60). (B) The effective haplotype number found in chromosomes I-II, III-IV, and V-VI (see Methods). Blue bars show the mean and one standard error of the mean. The loss of genetic diversity during experimental evolution implies that only a few genotypes could have been responsible for potential fitness responses due to randomized maternal effects.

